# GAT-HiC: Efficient Reconstruction of 3D Chromosome Structure via Residual Graph Attention Neural Networks

**DOI:** 10.1101/2025.04.18.649477

**Authors:** Beyza Kaya, Emre Sefer

## Abstract

Hi-C is an experimental technique to measure the genome-wide topological dynamics and three-dimensional (3D) shape of chromosomes indirectly via counting the number of interactions between distinct sets of loci. One can estimate the 3D shape of a chromosome over these indirect interaction datasets. Here, we come up with graph attention and residual network-based GAT-HiC to predict three-dimensional chromosome structure from Hi-C interactions. GAT-HiC is distinct from the existing 3D chromosome shape prediction approaches in a way that it can generalize to data that is different than train data. So, we can train GAT-HiC on one type of Hi-C interaction matrix and infer on a completely dissimilar interaction matrix. GAT-HiC combines the unsupervised vertex embedding method Node2vec with an attention-based graph neural network when predicting each genomic loci’s three-dimensional coordinates from Hi-C interaction matrix. We test the performance of our method across multiple Hi-C interaction datasets, where a trained model can be generalized across distinct cell populations, distinct restriction enzymes, and distinct Hi-C resolutions over human and mouse. GAT-HiC can reconstruct accurately in all these scenarios. Our method outperforms the existing approaches in terms of the accuracy of three-dimensional chromosome shape inference over interaction datasets. Code and datasets can be found at https://github.com/beyzoskaya/GAT-HiC.

## 1 Introduction

The chromosome’s three-dimensional (3D) structure has an impact on a number of genomic functions [1], [2], [3]. So, the discovery of chromosomes spatial 3D shape is crucial to understand regulatory and functional genome elements. To model the 3D shape of chromosomes and analyze each chromatin’s cellular and spatial organization, various chromosome conformation capture experiments were developed such as 3C [4], 4C [5], 5C [6], and Hi-C [7], [8], [9]. Overall, all chromosome conformation capture techniques focus on quantifying the number of contacts among genomic locations in order to understand the genome’s shape and organization. Among them, High-throughput Chromosome Conformation Capture (Hi-C) technique quantifies the number of interactions between the whole set of genomic locations at a whole genome level, where interactions are contacts between genomic location pairs in the genome. Hi-C experiment is composed of the following steps [7], [8], [9]: 1-A fixative solution is used to cross-link the chromatin among a number of chromosomes. 2-Chromatin is digested by a special enzyme after isolation. As a result of isolation and enzyme application, we obtain crosslinked DNA fragment pairs which are spatially close but their positions in the linear genome may differ remarkably. 3-DNA sequencing is used to amplify the templates which are formed after ligating the separate fragments again and reversing the crosslinks. 4-We obtain the ligation junctions frequencies at a genome level, where those relative interaction frequencies are associated with genomic positions spatial closeness in three-dimensional space. Hi-C is an all-to-all technique which allows us to analyze the entire genome’s global organization.

Hi-C technique has generated a large amount of interaction data which resulted in a number of computational techniques being developed. Those computational techniques focus on inferring chromosomes three-dimensional shapes by using their interaction data [10]. Many of these computational techniques employ a strategy involving distance-constrained optimization [11], [12], [13], [14], [15], [16]. In this case, these methods convert the input interaction matrix into distances via power law-based transformation [9]. These transformed distances are called Wish Distances. After converting the interaction frequencies into distances, three-dimensional x, y, and z coordinates are initially computed where each x, y, z coordinate combination maps to a chromosome location. Following such coordinate initialization, those x, y, and z coordinates are used to train the method via optimization such that the predicted three-dimensional structure’s Euclidean distances match with the input’s wish distances as accurately as possible.

Even though traditional distance-constrained approaches are frequently used, these traditional approaches have a number of limitations. First of all, several distance-constrained approaches make identical and independent distribution assumptions of the chromosome interactions [11]. However, this assumption is not valid since polymers self-attract which leads to a correlation among adjacent interaction locations [17]. Additionally, correlations between contacts are important knowledge so removing them will remove a key knowledge for three-dimensional shape prediction. Secondly, the existing distance-constrained approaches cannot be easily adapted across distinct problem instances. That means, when predicting the three-dimensional structure of a chromosome for a distinct interaction matrix, these methods train a method completely from scratch. In this case, distinct interaction matrices can be the matrices obtained over a different cell population, resolution, or restriction enzyme. As a result, when making predictions at higher resolutions, these methods require larger computational resources such as memory and CPU. Additionally, those conventional approaches do not perform well for sparser data since only a few attributes can be used during training.

Our GAT-HiC can store a trained model and predict the three-dimensional shape of unobserved data accurately by using such a trained model. Such ability to project knowledge is the definition of generalizability. Particularly, we will discuss generalizability over 3 dataset variations which are also discussed in [18]:

1. **Generalizability over Hi-C dataset resolutions:** We can train a model of a chromosome by using Hi-C interaction dataset at a given resolution, and then use the model to infer the same chromosome’s structure on a possibly distinct Hi-C interaction dataset with a different resolution accurately. So, we can train our method on lower resolutions, and infer three-dimensional shapes at higher resolutions. By following this approach, we do not need to train our method from scratch for each higher resolution which decreases the computational cost associated with such training.
2. **Generalizability over Hi-C restriction enzymes:** We can train a model of a chromosome by using Hi-C interaction dataset formed by using a given restriction enzyme, and then use the model to infer the same chromosome’s structure on a possibly distinct Hi-C interaction dataset formed by using a different restriction enzyme accurately. By following this approach, we do not need to train our method from scratch for each restriction enzyme which decreases the computational cost associated with such training.
3. **Generalizability over cell populations:** We can train a model of a chromosome by using Hi-C interaction dataset of a given cell population, and then use the model to infer the same chromosome’s structure on a possibly distinct Hi-C interaction dataset of a different cell population accurately. So, we can train our method on sparse interaction matrix having less interaction between chromosome positions, and infer three-dimensional shapes at dense interaction matrices.

Those generalizations bring our deep learning-based GAT-HiC to have several advantages that are missing in the competing approaches except [18]. The first generalization on resolutions is important since some approaches have high computational demand and such demand prevents them from being used in making three-dimensional inferences at higher-resolution datasets. This generalizability decreases the running time of GAT-HiC at higher resolution inference. The second generalization is important since it discusses the robustness of our method to biases as a result of selected restriction enzymes. It ensures that GAT-HiC performance is not impacted by the selected restriction enzyme in the training dataset. The last generalization exhibits the robustness of our GAT-HiC to possible sparser interaction datasets. In terms of validation, we evaluate the performance over 3 datasets of human (GM12878, K562) and mouse (mESC, neuron, and NPC) cell lines. We compared our performance with the best-performing inference methods: HiC-GNN [18], LorDG [11], and ShNeigh2 [19].

Here, we come up with a different type of distance-constrained approach GAT-HiC to reconstruct three-dimensional chromosomal structure over Hi-C interaction matrices, where our approach also handles the aforementioned disadvantages of conventional distance-constrained approaches. Our proposed solution is based on an attention-based and graph-based analysis of interaction datasets. In our graph-based analysis and development, we use the unsupervised vertex embedding approach Node2vec [20] in generating embedding attributes associated with each chromosome position. Afterward, we train a graph attention neural network (GAT) [21] with residual connections to use these unsupervised embedding attributes in predicting the three-dimensional coordinates of each chromosome position.

The paper is organized as follows: The related methods brief survey is provided in Section 1.1. Section 2 will give details about GAT-HiC, its unsupervised embedding component as well as residual graph attention neural network. The details about the experimental evaluation are described in Section 3. Finally, our results are discussed in Section 4, and the conclusion is discussed in Section 5.

### 1.1 Related Work

A number of approaches have been proposed to reconstruct three-dimensional chromosome shapes. One of them is MCMC5 where Markov Chain Monte Carlo (MCMC) is used to sample three-dimensional coordinates over the posterior distribution. Such posterior distribution uses a normal distribution prior over the interaction frequencies [22]. Another method BACH is similar to MCMC5 except the distribution is assumed to Poisson instead of normal distribution [23]. Similarly, PASTIS makes a Poisson distribution assumption for the relationship between three-dimensional coordinates, distance, and interaction frequencies. But PASTIS maximizes the likelihood in its optimization [24]. Chromosome3D, as a distance-constrained approach, uses simulated annealing to optimize the corresponding distances [15]. LorDG [11] is also a distance-constrained approach which optimizes a Lorentzian function in its objective. In this case, the Lorentzian objective function focuses on smoothing inconsistent Hi-C data which may be a result of a population of heterogeneous cells. Moreover, the Lorentzian objective function rewards the consistent constraints fulfillment. ChromSDE [16], as a distance-constrained approach, utilizes Semidefinite Programming to infer three-dimensional structures. When inferring the relation function between distance and contact frequencies, ChromSDE uses a golden search method. Among these distance-constrained approaches, we will be comparing GAT-HiC with ChromSDE and LorDG since they perform better than a number of remaining contact-based approaches which directly utilize interaction data in reconstructing three-dimensional structures [10], [11], [16], [25]. These approaches are the top-performing ones so they can represent contact-based approaches in reconstructing three-dimensional structures.

Among other categories of methods, we also evaluate the performance GAT-HiC with respect to ShRec3D and ShNeigh2. These two approaches take the Hi-C interaction graph’s neighbourhood structure into account so they can be considered as graph-based approaches similar to GAT-HiC. Due to their inherent difference from the methods discussed above, we include these approaches in our comparison. Among them, ShRec3D [26] runs the shortest path algorithm over the interaction matrix dataset as part of distance derivation from interactions. ShRec3D analyzes the interaction locations neighbourhood by using a shortest path algorithm. Afterward, it applies multidimensional scaling to infer three-dimensional structure from these calculated distances. On the other hand, ShNeigh [19] is another approach which defines an affinity matrix to take neighbourhood into account. This affinity matrix is related to the input interaction matrix, and its entries are added to the objective function as regularization terms during optimization. ShNeigh comes with 2 versions; ShNeigh1 and ShNeigh2 where the difference is their assumption of the relationship between contact frequency and distance. ShNeigh2 treats this relationship as dynamic and optimizes it, whereas ShNeigh1 assumes a static dependency between contact frequency and distance. In our case, we compare GAT-HiC with ShNeigh2 since it generally results in a better performance.

The most similar approach to ours is HiC-GNN [18] which uses a convolutional graph neural network to infer three-dimensional chromosome structures. Their convolutional neural network structure could take the Hi-C interaction graph’s neighbourhood structure into account so it can also be considered a graph-based approach. However, our attention-based graph neural network with its residual connections is superior to the convolutional graph neural network as well as our Node2vec-based unsupervised embedding approach being more capable.

## 2 Methods

GAT-HiC uses a contact map as an input for each chromosome. Contact map contains the interaction frequencies between different genomic loci of the chromosomes. For a given chromosome, each row represents the interaction frequency between two different genomic loci. The contact map is represented by *N × N* symmetric matrix *CF*, where *N* represents the number of genomic loci. Each row of the contact map matrix is used to create a weighted graph with *N* nodes, which are assumed to be adjacent to each other. If there is no weight (interaction) between two loci, the two loci mentioned are not connected. In this way, predicting the structure of the chromosome via GAT-HiC can be interpreted as a regression of the nodes in the graph.

### 2.1 Conversion of Contacts to Wish Distances

The main problem in three-dimensional structure prediction is that there is no ground truth for the input matrix. In general, our goal is to find output coordinates via our approach such that true distances between input chromosome locations match our estimated distances. However, we do not know the true distances between input chromosome locations. Empirical and theoretical studies discuss there is an inverse exponential relation between interaction count between two locations and the corresponding distance between the same locations [9], [25], [27], [28]. According to these studies, let *CF*_*ij*_ be the total interaction count between segments *i* and *j*, true distance between segment *i* and *j* can then be calculated as:

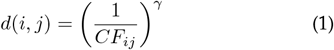

where *γ* defines the conversion factor. Generally, *γ* is not known and its value is impacted by the considered chromosome. For many cell types, *γ* is between 0.1 and 2 [29]. As part of the experiments, we fix the conversion factor to be 0.1.

### 2.2 Node Embedding Creation

Reconstructing the 3D structure of chromosomes from Hi-C contact maps requires translating the interaction frequencies into a form that can be effectively utilized by machine learning models. Since Hi-C data solely shows weighted connections between points, relating to how often genomic locations interact, there are no built-in characteristics for the points in the graph. Creating significant node representations is, therefore, an essential stage in preparing the data for subsequent tasks. These representations ought to accurately reflect the structural property of the contact map while offering a definition that helps in predicting 3D spatial connections between locations.

To tackle this, we use Node2vec [20] algorithm to produce node representations. Node2vec is an adaptable and expandable method for node representation that captures both local and global graph forms by modeling biased random walks on the graph. Unlike other representation techniques, Node2vec provides the ability to manage the balance between breadth-first search (BFS) and depth-first search (DFS) movement through two hyperparameters, *p* and *q*. This adaptability allows the method to fit the hierarchical and local connection patterns found in chromosomal contact maps, leading to representations that accurately reflect the weighted graph form of Hi-C data.

By optimizing the probability of detecting a node’s context (which is its neighbors in the random walk), Node2vec algorithm learns embeddings. In particular, the goal for the two genetic loci *i* and *j* can be expressed as follows:

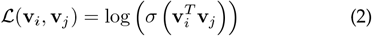

where ℒ (**v**_*i*_, **v**_*j*_) is the objective function that maximizes the likelihood of nodes *i* and *j* interacting. Here, **v**_*i*_ and **v**_*j*_ represent the embedding vectors for nodes *i* and *j*, respectively, while 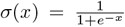 is the sigmoid function that converts the dot product 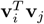 (the similarity between nodes *i* and *j*) into a probability score between 0 and 1. The objective function maximizes the likelihood of identifying node *j* in relation to node *i*. The probability that nodes *i* and *j* are interacting in the graph is determined by the dot product of the embedding vectors **v**_*i*_ and **v**_*j*_, which are learned by Node2vec using random walks.

By learning embeddings that reflect the combination of loci in random walks, Node2vec captures the structural connections between loci in the context of Hi-C data, where nodes stand for genomic loci and edges for interaction frequencies. The bias can be adjusted by the parameters *p* and *q*, which modify the ratio of depth-first to breadth-first search methods. This is accomplished by conducting biased random walks across the loci graph. In subsequent tasks, such as 3D genome structure prediction, where the spatial relationships between genomic loci are deduced from their embeddings, the inferred embeddings can be used.

### 2.2.1 Hyperparameter Optimization for Node2vec

We performed a grid search over multiple Node2vec hyper-parameter combinations to find the optimal settings for our application in order to optimize the embeddings for GAT-HiC. The following hyperparameters were examined, along with their ranges:

- **Embedding dimension**: 128, 256 and 512.
- **Walk length**: 50 to 250 steps.
- **Number of walks per node**: 20 to 50.
- **Window size**: 5 to 30.
- **Return parameter (***p*): 0.25 to 4.
- **In-out parameter (***q*): 0.25 to 4.

### 2.3 Hi-C Map Normalization

Graph neural network takes two inputs: 1-The attributes of nodes, 2-Associated Hi-C interaction matrix. In this case, the Hi-C interaction matrix is an adjacency matrix which corresponds to a weighted graph. Such weighted graph’s weights model the matrix’s interaction count. Those interaction count values are frequently in the order of a hundred thousand. In order to make the graph neural network numerically stable, the input interaction matrix and associated weights are normalized into [0, 1] interval. In our case, we use Knight-Ruiz (KR) matrix balancing to normalize our matrix [30]. KR balancing has frequently been utilized in a number of Hi-C datasets [31]. KR balancing generates a doubly stochastic matrix.

### 2.4 Graph Attention Selective Residual Network

To develop complex node representations from graph data, GAT-HiC combines layer normalization, residual connections, and Graph Attention Networks (GAT) [21]. For tasks such as regression, as 3D coordinate prediction in our case, feature learning is essential, and each layer of the network contributes significantly to its improvement.

In graph-based data, where node interactions are crucial, graph attention networks are particularly well-suited []. In contrast to conventional graph convolutional networks (GCNs), GATs use a self-attention mechanism that enables the model to dynamically assess the significance of adjacent nodes. This allows the network to concentrate on the most relevant neighbors rather than treating every node equally, which is particularly crucial when working with sparse and large-scale graph structures. Figure 1 summarizes GAT-HiC pipeline.

**Figure 1:**
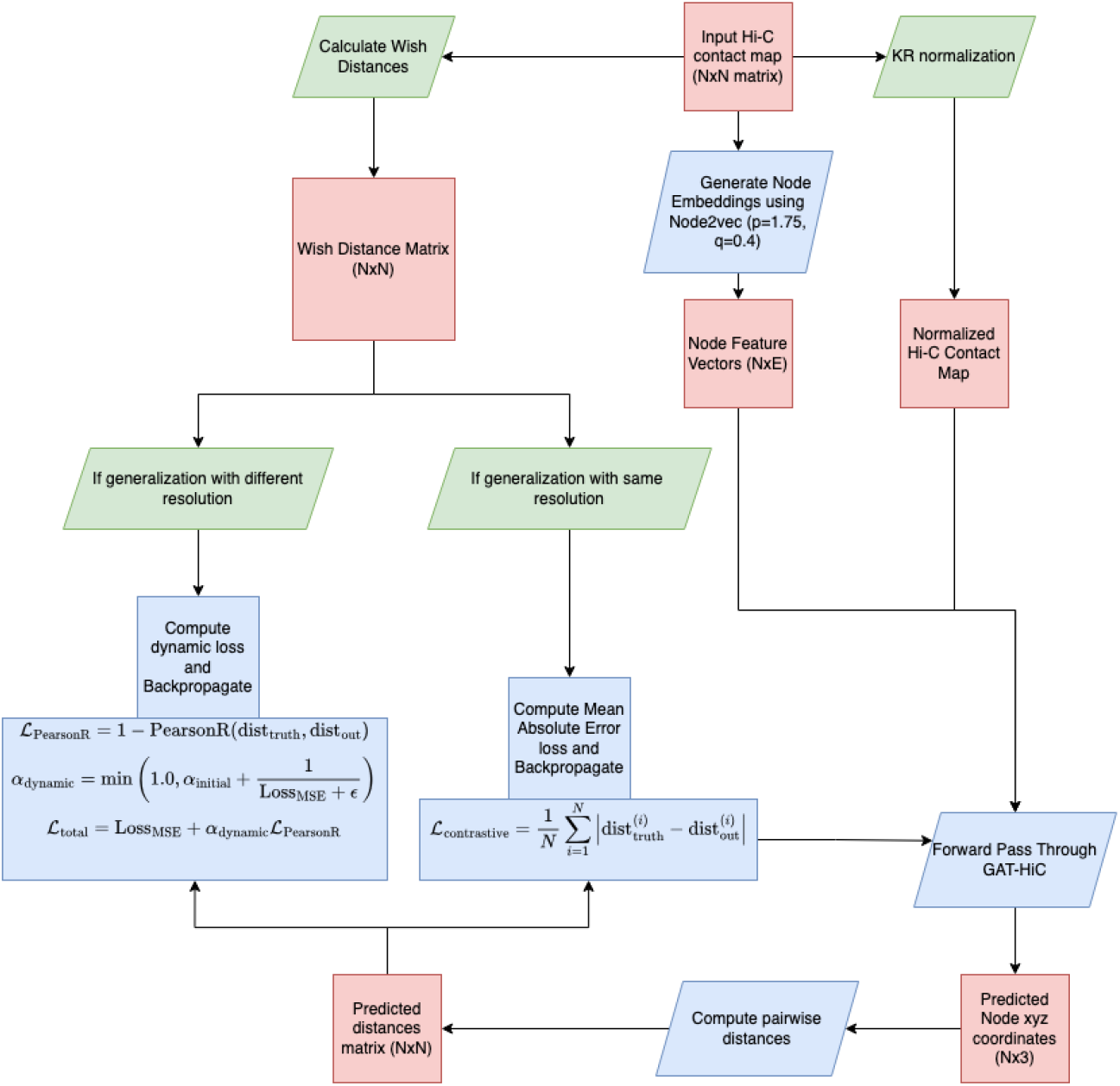
GAT-HiC starts with raw Hi-C data, generating node embeddings via Node2Vec. After processing with GATv2Conv [32] layers, dense layers with residual connections and layer normalization refine the feature vectors. The model predicts 3D spatial positions by calculating pairwise distances between node embeddings, optimized using task-specific losses: contrastive loss for same-resolution data and a combination of dynamic weighted Pearson loss with MSE for generalization.

Our model’s GATv2Conv [32] layer utilizes the feature vectors of the target node *i* and its neighbors *j* to compute attention scores *α*_*ij*_, which aggregate information from adjacent nodes. For graph data with varying node importance, such as genomic or protein interaction data, these attention coefficients enable the model to concentrate more on significant neighbors throughout the feature aggregation phase. This layer assists the model in producing more precise node embeddings by capturing intricate dependencies in the network. The revised node characteristics are calculated formally as follows:

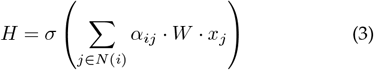

where **W** is a learnable weight matrix, *α*_*ij*_ is the attention weight between nodes *i* and *j, N* (*i*) denotes the neighborhood of node *i*, **H** represents the updated node characteristics, and the nonlinearity *σ* is the ReLU activation function. We include residual connections in the denser parts of the model to improve learning and ensure network stability. By enabling gradients to avoid one or more layers, residual connections are an effective way of mitigating the problem of vanishing gradients that can arise in deep networks. This is especially crucial when deep networks are being trained on graph data, as learning high-level representations while maintaining low-level feature information is vital. GAT-HiC incorporates a residual connection to merge the output with the original features following each dense layer transformation. Particularly, the feature representations at each layer *l* are updated by complementing the transformed features *f* (**H**^(*l*)^), where *f* (·) represents the transformation applied to the features at layer *l*, composed of a linear transformation followed by the ReLU activation function, with the original input features **H**^(*l*)^. This ensures that essential details from earlier layers are retained, helping the model preserve important low-level features while learning complex representations through deeper layers. The updated feature representation is computed as:

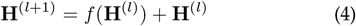

A smoother gradient flow during backpropagation is achievable by the inclusion of the original input features to the transformed features. This enhances training efficiency both in global and local structures for the chromosomes and helps prevent overfitting, particularly in complicated problems like 3D structure prediction.

We use layer normalization following each dense layer to further stabilize the training process and enhance convergence. By normalizing the activations to have a zero mean and unit variance, layer normalization aids in standardizing the activations across the features. This keeps any feature from affecting the training process and ensures that the learning rate remains constant across the network. For each layer, normalization is done using the following equation:

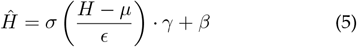

where *ϵ* is a small constant to avoid division by zero, *γ* and *β* are learnable parameters, and *µ* and *σ* are the features’ mean and standard deviation, respectively. A ReLU activation is then applied to the normalized features 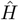.

The network uses a number of dense layers after the first GATv2Conv layer to decrease the node feature dimensions while maintaining structural information that is essential for 3D coordinate prediction. Normalization and a ReLU activation come after each layer, preventing overfitting while preserving useful node representations. With fewer parameters, this dimensionality reduction increases model efficiency.

The model applies a distance metric to the node features after it has passed the dense layers in order to calculate the final output. In particular, pairwise Euclidean distances between the final embeddings of every node are computed. The similarity between the expected and actual places is assessed using these distances, which is essential for tasks involving the prediction of 3D structures.

Two distinct loss functions are used by the model, depending on the type of data. A contrastive loss is applied for data with the same resolution, and a dynamically weighted Pearson loss combined with Mean Squared Error (MSE) is utilized for generalization across resolutions. The following subsections will provide a more thorough explanation of these losses. For a better understanding of the model’s overall flow, the complete GAT-HiC workflow is illustrated in Figure 1, while Figure 2 shows the detailed architecture of the model. Our neural network parameters are optimized via backpropagation by using Adam optimizer [33], where the mean squared error between the output structure’s distances and wish distances is minimized.

**Figure 2:**
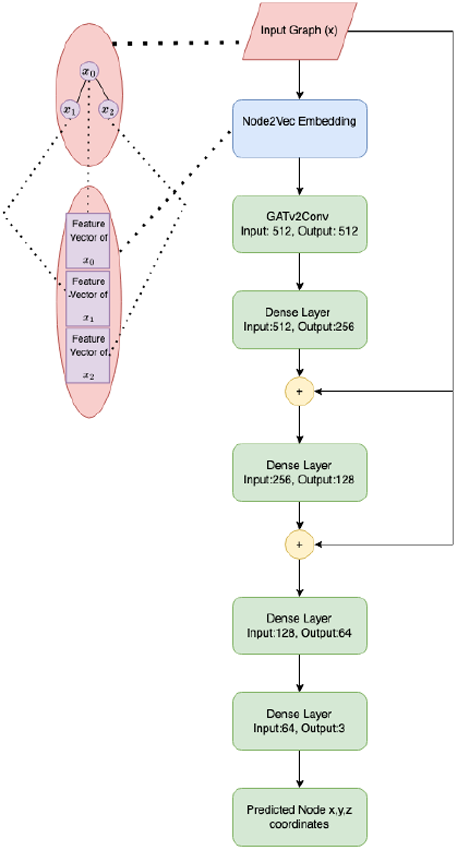
Residual Graph Attention Neural Network architecture diagram. This figure illustrates specific layers in the model.

### 2.5 Task Specific Loss Functions

While predicting xyz coordinates via contact maps, predictions are made on data with the same resolution, as well as generalizations are made between different resolutions.

While it is important to represent global information while making generalizations between different resolutions, it is more important and necessary to capture local information for data with the same resolution. When we used Mean Square Error as a loss function, we could not get good results because it converged very quickly and we designed two different loss functions for two different tasks.

#### 2.5.1 Generalization Between Different Resolutions Loss Function

We created an integrated loss function that establishes a balance between local quality and global consistency in order to enable consistent generalization across datasets with different resolutions. This is accomplished by combining a dynamically weighted Pearson correlation loss with the Mean Squared Error (MSE) loss. This allows the model to retain large-scale structural interactions within the predicted 3D chromosomal structure while also capturing fine-grained details.

Accurate predictions of individual distances are provided by the MSE loss, which calculates the element-wise differences between the predicted and true pairwise distances. The global structural consistency of the 3D structure, however, may be lost if MSE is the only method used, particularly when extending over datasets with different resolutions. We use a Pearson correlation loss, which calculates the correlation between the true and predicted distances, to address this limitation. The Pearson correlation coefficient, *ρ*, is defined as:

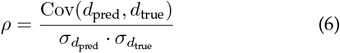

where *d*_pred_ and *d*_true_ represent the predicted and true distances, Cov denotes covariance, and *σ* represents the standard deviation. The Pearson correlation loss is then expressed to overcome the minus loss in each iteration:

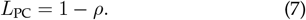

In order to maintain global dependencies in the 3D structure, *L*_PC_ term encourages the model to optimize the linear agreement between predicted and true distances. When combining MSE loss with *L*_PC_, we introduce a dynamic weighting factor, *α*_dynamic_, to handle the relative importance of both losses during training in order to balance their contributions. The weighting factor is defined as:

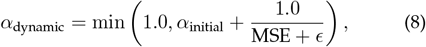

where *ϵ* is a small integer to prevent division by zero and *α*_initial_ is a small constant (e.g., 0.1). This formulation emphasizes global consistency in the later training stages by allowing the Pearson correlation loss to have a larger impact as the MSE decreases. The final loss function is given by:

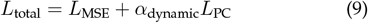

To ensure local accuracy, the initial term of the final loss places a significant value on making precise pairwise distance predictions. In order to promote overall consistency throughout the 3D structure, the second term concentrates on preserving global structural interactions.

#### 2.5.2 Same Resolution Loss Function

Local information is crucial to obtain precise and highquality 3D structure predictions for same-resolution data, when the structures are expected to be captured with fine detail. This is due to the fact that the model needs to focus on specific details and small changes in the distance relationships between loci while maintaining a high level of resolution. Standard loss functions, such as MSE, may not be adequate in this situation to capture these complex local dependencies, which would result in a loss of resolution in the expressed structures.

To address this, we use the mean absolute error loss (L1 loss), which is more sensitive to the local variations between the true and predicted distances. This loss makes sure that the model learns more effectively to retain the relative distances between the nodes by concentrating on the absolute difference between predicted and true pairwise distances. In same-resolution data, where fine-grained precision directly affects the performance metrics such as dSCC (domain-specific correlation coefficient), this is essential for preserving the high fidelity needed. Our loss is mathematically defined as:

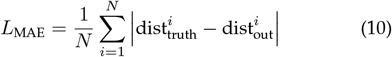

where dist_truth,*i*_ is the true pairwise distance between the nodes, dist_out,*i*_ is the predicted pairwise distance for the same nodes, and *N* represents the number of node pairs considered. By utilizing the L1 loss, the model captures intricate local relationships that are crucial for the same-resolution data, effectively emphasizing the fine-grained differences between the true and predicted distances.

### 3 Experimental Details

### 3.1 Evaluation

In grid search, we focused on optimizing vertex embedding parameters, graph neural network’s hidden layers sizes, the weighting factor, and learning rate. We evaluated the reconstruction performance via distance Spearman Correlation Coefficient (dSCC) which is a non-parametric metric using rank correlation. Generally, dSCC values close to 1 indicate a better performance. dSCC is defined as:

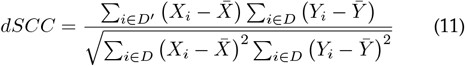

where pairwise distances between model’s all loci are represented by *D*^*′*^. Among the remaining ones, *X*_*i*_ defines the rank of distance *i* in *D*^*′*^, *D* represents wish distances of the chromosome, and *Y*_*i*_ is the rank of wish distance *i* in *D*. 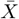 and 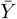 define the average of ranked vectors in *D*^*′*^ and *D* respectively.

dSCC defines a rank correlation which can evaluate the inference performance in a scale-invariant way unlike mean squared error. For instance, GAT-HiC might find an exact matching for chromosome, but associated three-dimensional coordinates might be scaled by a constant. If we use a non-ranking correlation measure, such a constant will have a decreasing effect. Such a decrease in non-ranked correlation is wrong since it only implies dissimilarity between inferred structure and ground truth in terms of just scale and location. The aim of modeling chromosomes in three-dimensional space is for visualization, so output’s scale is not important. As a result, dSCC is a valid structural similarity metric in our case. Our code and datasets can be found at https://github.com/beyzoskaya/GAT-HiC.

### 3.2 Real Hi-C Data

We test three-dimensional reconstruction performance over 5 real Hi-C datasets corresponding to human and mouse cell lines. In GM12878 cell line dataset [34], the dataset is formed by using Mbol restriction enzyme at 250 kb, 500 kb, and 1 Mb resolutions over 23 chromosomes. This dataset is utilized in testing the first generalization. To test the second generalization, we use mouse mESC, neuron, and NPC cell lines [35] by using Ncol and HindIII restriction enzymes over 22 chromosomes at 1 Mb resolution. Lastly, in K562 cell line dataset [34], we have a number of Hi-C interaction matrices over 23 chromosomes at 1 Mb resolution. This dataset is utilized in testing the third generalization.

## 4 Results

### 4.1 Generalizability Over Dataset Resolutions

To evaluate the performance of GAT-HiC across different resolutions, we used a GM12878 cell line dataset formed via the Mbol restriction enzyme with three resolutions: training was performed at 1 mb resolution, and testing was conducted at 500 kb and 250 kb resolutions. To combine embeddings from different resolutions, we applied the alignment method from HiC-GNN. With aligned embeddings, we assessed performance using the distance Spearman Correlation (dSCC) on the higher-resolution data. To optimize performance for each chromosome, we performed a grid search to select different hyperparameters adjusted to the unique structural characteristics of each chromosome.

To assess GAT-HiC’s ability to establish correlations across different resolutions, we first trained the model using 1 mb resolution contact maps along with low-resolution data. Afterward, the embeddings generated for 500 kb and 250 kb resolutions were aligned with the 1 Mb embeddings. The trained model was then tested on 500 kb and 250 kb resolution data. Since different chromosomes exhibit varying structural characteristics, hyperparameters such as the learning rate and convergence threshold were adjusted individually for each chromosome. The learning rate was chosen within the range of 0.0005 to 0.001, while the convergence threshold was selected between 1 × 10^*−*6^ and 1×10^*−*8^ based on grid search results tailored to each chromosome’s specific structure.

The model performance across different resolutions was compared by evaluating the dSCC values for the HiC-GNN model, which was trained at the 1 Mb resolution and tested at the 500 kb and 250 kb resolutions. These results were then compared with the dSCC values obtained using GAT-HiC. As shown in Figure 3, GAT-HiC outperformed HiC-GNN in terms of dSCC for most chromosomes at the 500 kb and 250 kb resolutions. However, due to the varying structures and higher variance in some chromosomes, GAT-HiC achieved lower dSCC values for chromosomes with greater variance. Figure 4 provides a detailed view of the chromosomes exhibiting high variance in the performance between different resolutions.

**Figure 3:**
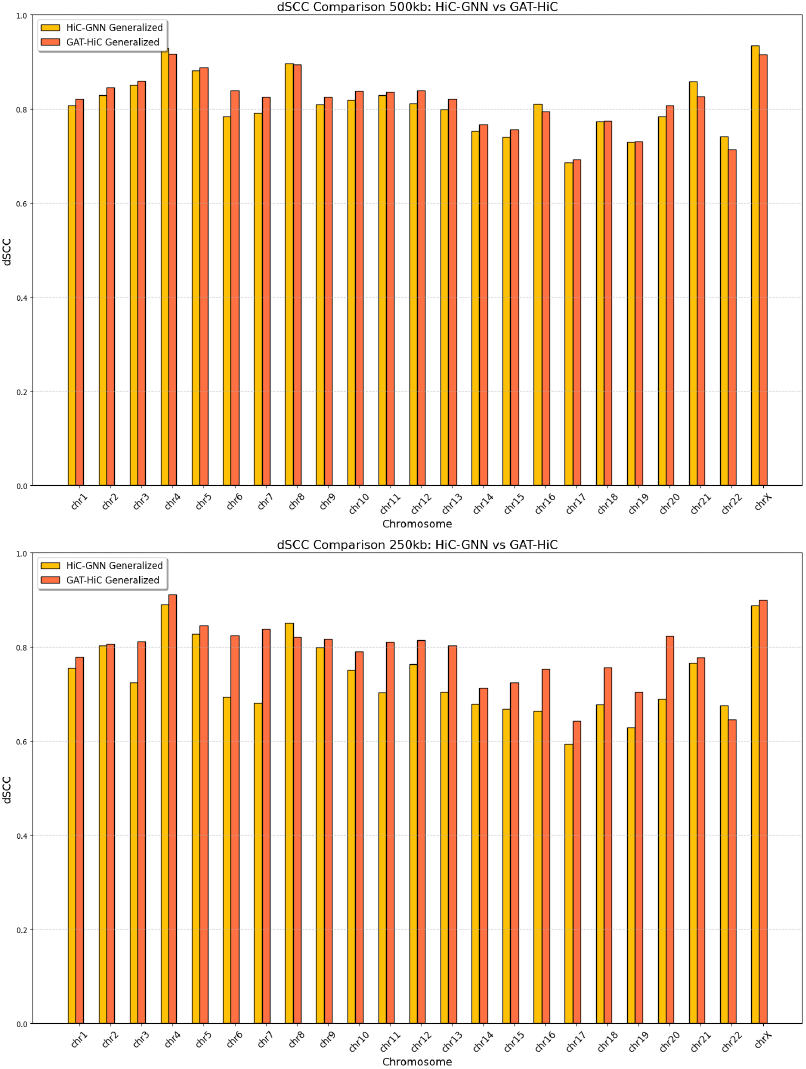
dSCC values between different resolutions. The dSCC values for HiC-GNN and GAT-HiC models are compared. The models are trained at 1 mb resolution and tested at 500 kb and 250 kb resolutions.

**Figure 4:**
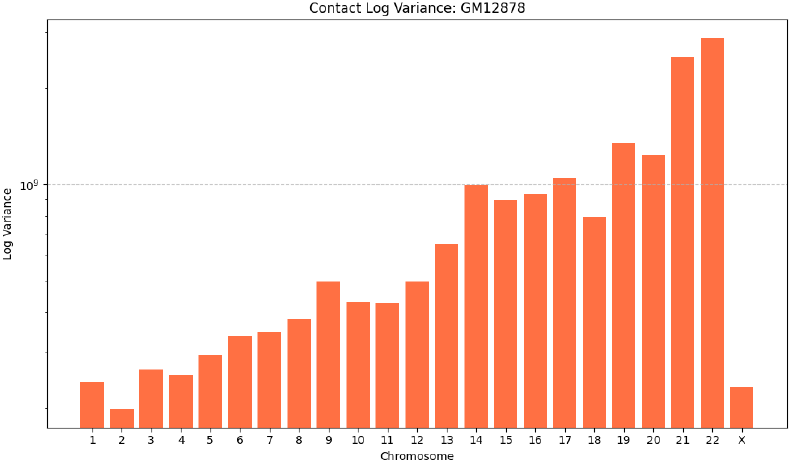
The contact log variances for GM12878 dataset’s each chromosome.

The performance of GAT-HiC was evaluated by comparing its dSCC values against three existing methods: HiC-GNN [18], ShNeigh2 [19], and LorDG [11]. These models were chosen because of their established effectiveness in chromosomal structure prediction and their ability to handle Hi-C data across various resolutions. HiC-GNN is a graph-based approach adjusted for Hi-C data, making it a direct alternative method for comparison. ShNeigh2 utilizes spatial networks, and LorDG is a deep learning model combining graph representations with resolution scaling. The models were evaluated at three different resolutions: 1 mb, 500 kb, and 250 kb. The results show that GAT-HiC consistently outperforms all three methods across most chromosomes, achieving higher dSCC values. This is particularly true for the 500 kb and 250 kb resolutions, where GAT-HiC performs better in capturing both local and global relationships, resulting in higher accuracy even for chromosomes with complex structural variations. These findings demonstrate that GAT-HiC offers superior performance in 3D chromosomal structure prediction compared to the existing methods. Figure 5 illustrates the dSCC values for each model at 1 mb, 500 kb, and 250 kb resolutions, where our GAT-HiC achieves superior performance compared to the other methods.

**Figure 5:**
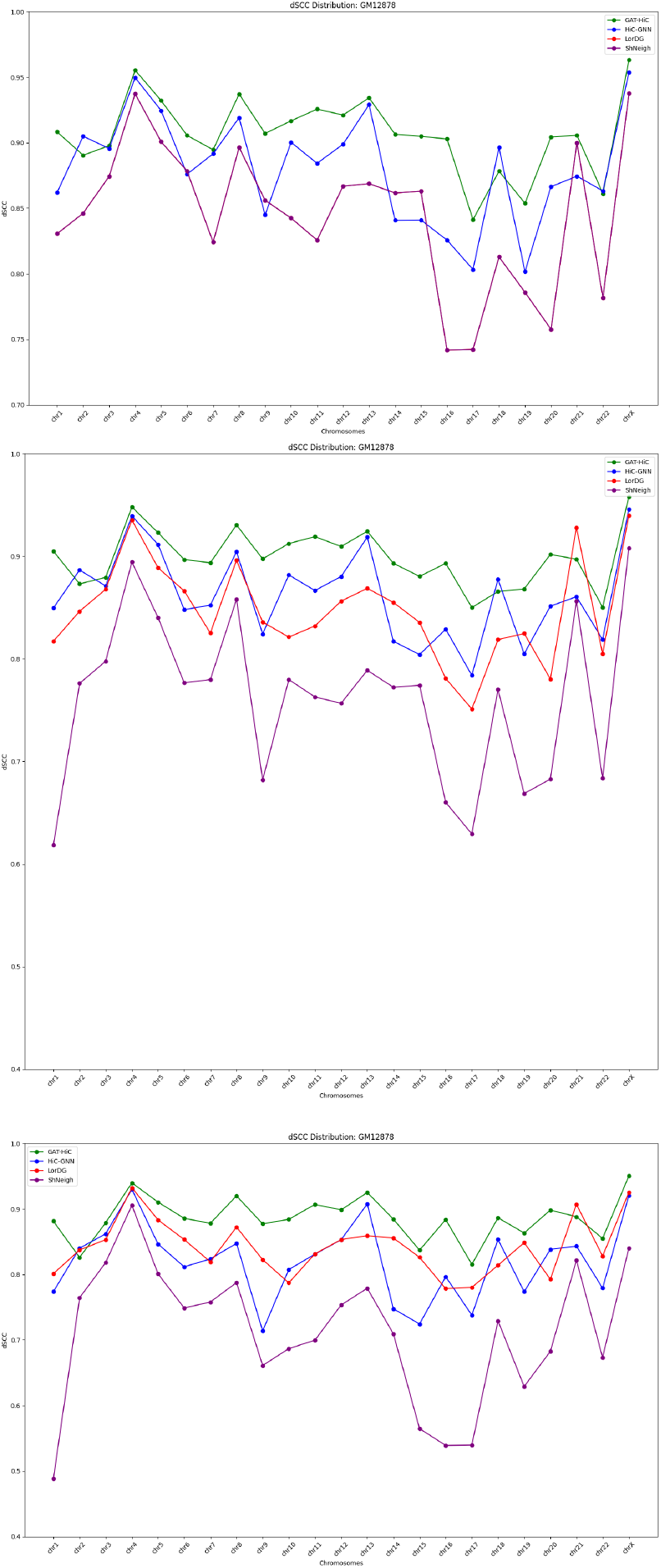
Comparison of dSCC values between different methods across various resolutions (1 mb, 500 kb, and 250 kb). The figure shows the comparison of GAT-HiC with other methods on GM12878 data. GAT-HiC achieved significantly higher dSCC values compared to other methods for most chromosomes.

### 4.2 Generalizability over Restriction Enzymes

Evaluating the generalizability of GAT-HiC under varying experimental conditions required the use of datasets generated with two different restriction enzymes: NcoI and HindIII. We tested the generalizability of restriction enzymes on mouse mESC, neuron, and NPC cell types [35].

In this dataset, we have 20 chromosomes interaction maps obtained via HindIII and Mbol restriction enzymes over 1 Mb resolution. These enzymes were chosen due to their widespread use in chromatin conformation studies and their ability to capture distinct aspects of genomic interactions. Restriction enzyme has an impact on the output interaction matrix which varies from enzyme to enzyme [36], [37], [38]. So, generalizability over restriction enzymes will show the robustness of GAT-HiC to this variation. We test this scenario by training method on one restriction enzyme, and test the performance on another enzyme.

We compare the performance of GAT-HiC with HiC-GNN over datasets generated by using two different restriction enzymes in terms of dSCC. This comparison highlights the adaptability of GAT-HiC to varying experimental conditions and its ability to capture chromosomal interaction patterns effectively. Figure 6 represents the dSCC values for each chromosome across both NcoI and HindIII enzymes. Notably, GAT-HiC outperforms HiC-GNN on nearly all chromosomes, demonstrating its strong performance in accurately preserving structural information.

**Figure 6:**
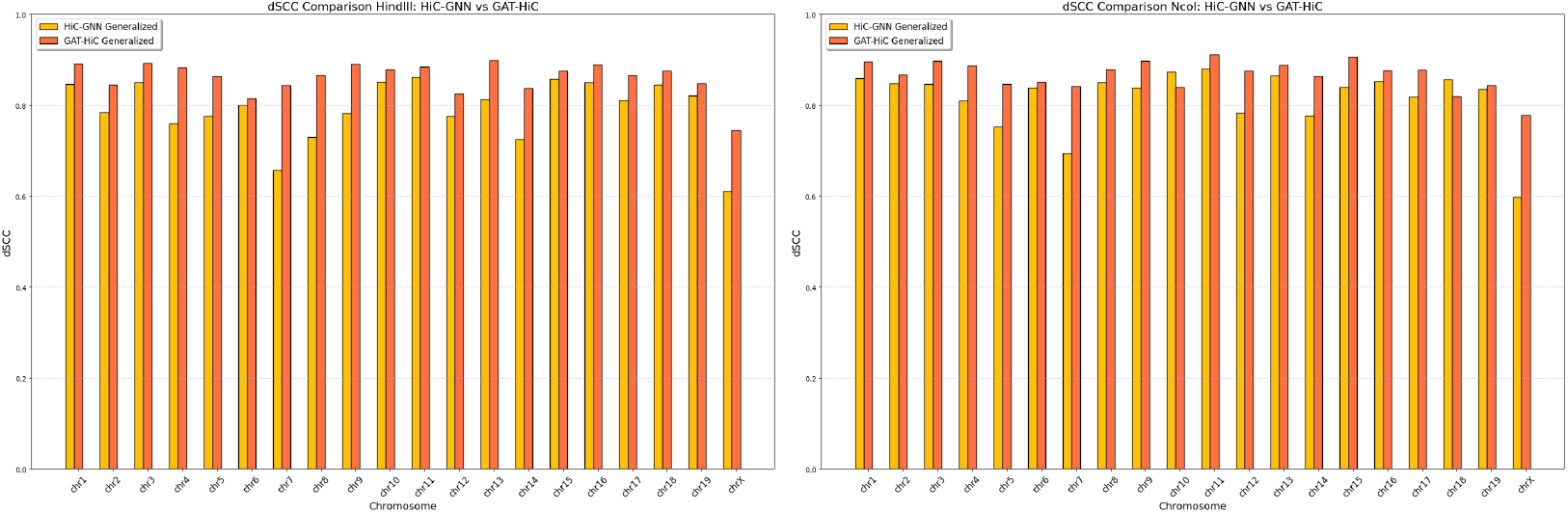
Comparison of dSCC values between GAT-HiC and HiC-GNN on Hindlll and Ncol enzymes.

The performance of the models is further demonstrated by the distance comparison graphs for chromosomes 5 and 12 in Figure 7, which builds on the comparison of GAT-HiC with HiC-GNN across datasets generated with two different restriction enzymes. Despite the apparent similarity in the predicted distances for the two approaches, there are minor but notable variations in the global structures. These variations help explain why GAT-HiC has higher dSCC values, indicating that it is more effective in capturing structural links between loci. The improved performance of GAT-HiC depends critically on the local structural reliability, even when the overall distance trends are similar. This highlights how crucial it is to adjust local structural characteristics because minor modifications can have a significant effect on dSCC values.

**Figure 7:**
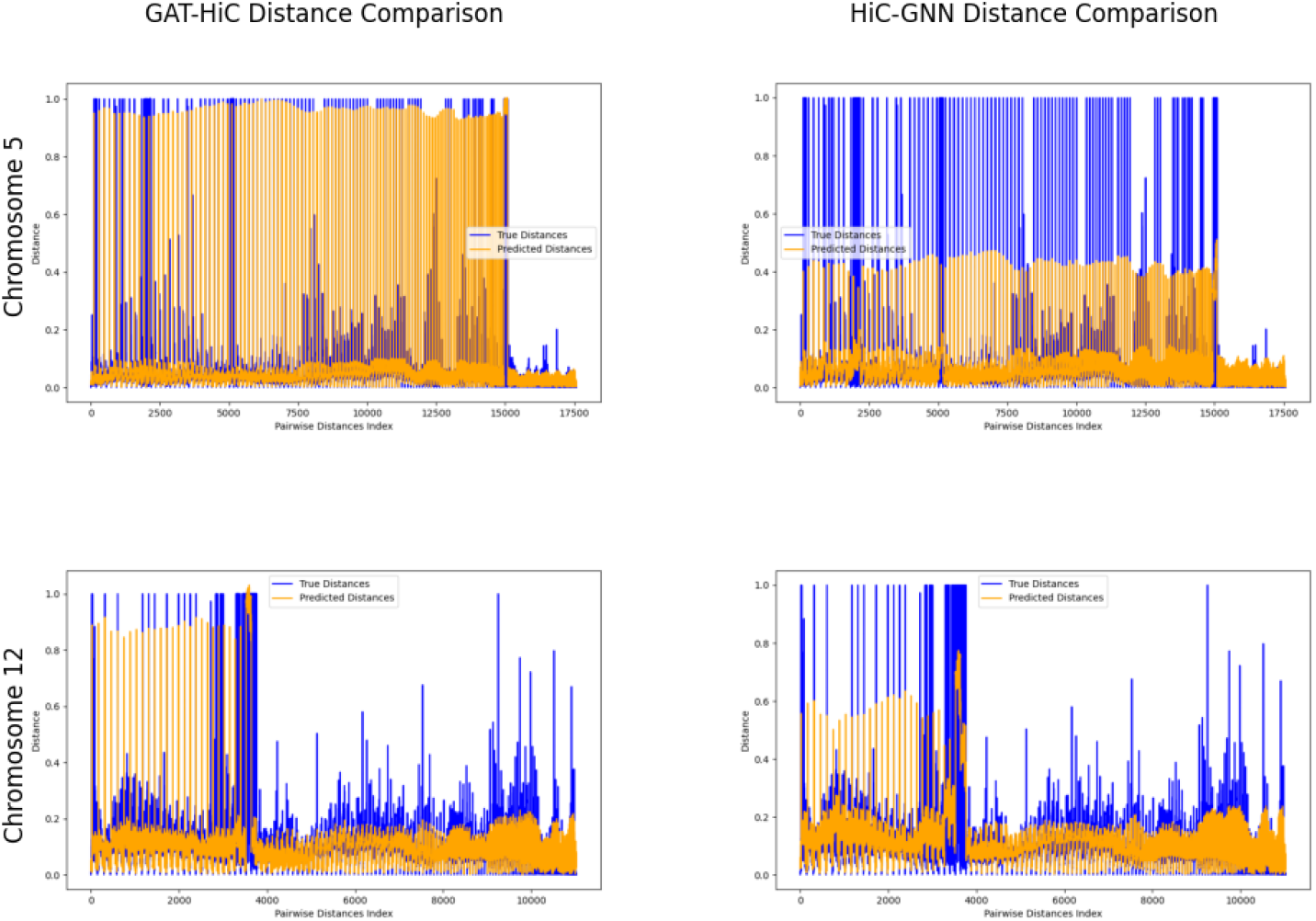
The distance comparison between predicted and ground truth. The first column shows the GAT-HiC distance comparison for chromosome 5 and chromosome 12 and the second column shows the distance comparison for HiC-GNN for same chromosomes.

### 4.3 Generalizability over Cell Populations

We evaluate how well GAT-HiC performs on two datasets—full and half coverage data that are obtained from the K562 cell line. We utilized K562 cell line from [34] where the dataset is obtained over 22 autosomal chromosomes at 1 mb resolution which are obtained over several replicate experiments. Each replicate is generated over distinct cell populations. Each replicate has different interaction counts since each replicate is different in population size. Full coverage datasets provide an extensive comprehension of chromosomal interactions with interaction data available over the entire chromosome. Half coverage datasets, on the other hand, are condensed versions of the data that only take into account a subset of interactions, usually encompassing approximately half of the chromosomal locations. By assessing GAT-HiC on both coverage levels, we aim to demonstrate how effectively the model works with varied data sparsities and how efficiently it can identify patterns of chromosomal interactions under different conditions.

Figure 8 illustrates the dSCC values for GAT-HiC and HiC-GNN on both half and full coverage data, showing the model’s performance across 22 chromosomes. As seen in the figure, GAT-HiC consistently outperforms HiC-GNN in terms of dSCC values for both datasets, indicating its robustness in capturing interaction patterns, even with reduced coverage.

**Figure 8:**
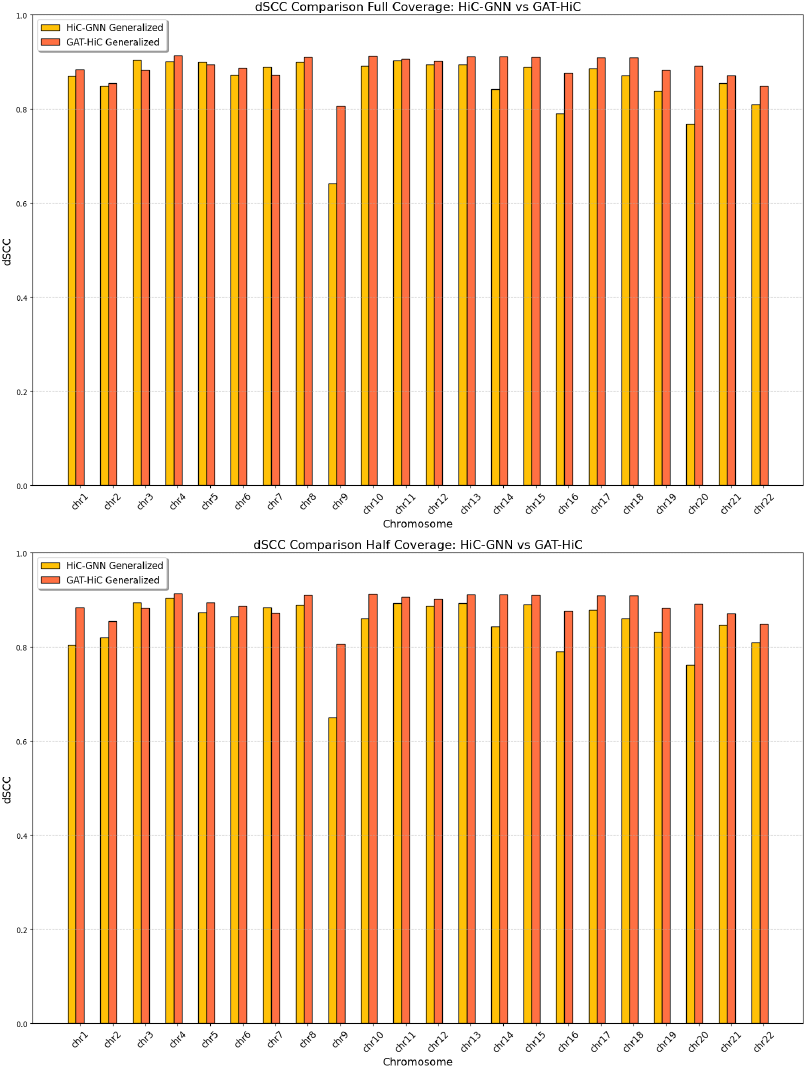
Comparison of dSCC values between GAT-HiC and HiC-GNN on Half and Full coverage on 22 chromosomes in K562 data cell.

Generalization across different cell populations was explored further by training our GAT-HiC on the half-coverage dataset and testing its performance on the full-coverage dataset. This case assesses the model’s ability to adapt from sparse data to denser data, a common challenge in Hi-C experiments. As illustrated in Figure 9, we compared the dSCC values between HiC-GNN and GAT-HiC across all chromosomes. The results reveal that GAT-HiC consistently outperforms HiC-GNN in most chromosomes, achieving higher dSCC values. This improvement highlights the robustness of GAT-HiC in addressing the transition from sparse to dense Hi-C data. The sparse half-coverage dataset introduces missing interactions, making it challenging to infer chromatin structures accurately. However, GAT-HiC captures meaningful structural patterns more effectively, enabling it to generalize better when applied to full-coverage data. Figure 9 provides a detailed comparison, showing that GAT-HiC serves enhanced dSCC values across several chromosomes, reflecting its improved ability to maintain chromatin contact patterns under varying data sparsity conditions. Even though there is a reconstruction performance decrease when we generalize on data with fewer interactions, we note that generalized models were either trained or tested on datasets with half coverage.

**Figure 9:**
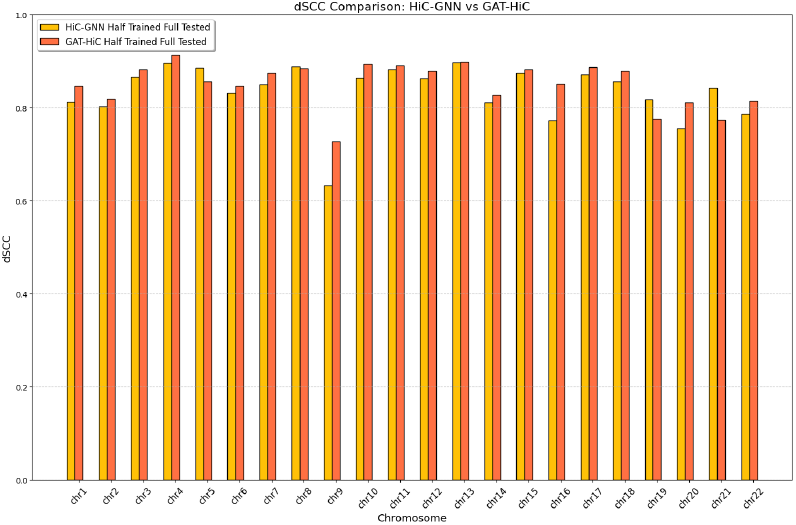
dSCC comparison between HiC-GNN and GAT-HiC for training on half coverage data and tested on full coverage.

## 5 Conclusion

Here, we come up with a novel approach GAT-HiC to predict three-dimensional chromosome structure by using Hi-C dataset. Our approach combines vertex embeddings with graph neural networks while making such predictions. Our inference approach is capable of generalizing across cell populations, restriction enzymes, and resolutions. Resultingly, our approach performs better than the existing methods over multiple datasets.

Our approach GAT-HiC could be generalized for 3 main reasons. First of all, we create static vertex attributes for each loci before training so the neural network’s trained parameters could be stored to infer over unseen data. To our best knowledge, all remaining structure inference approaches except one of them treat each location’s coordinates as trainable parameters which makes it impossible to integrate the trained parameters for structure prediction on a novel dataset. Second of all, the vertex attributes that are created are alike among multiple datasets in order for the following neural network results to be compatible. We have an increased prediction performance once we align embeddings, so vertex embeddings of Hi-C interaction matrices are roughly isometric. Due to this isometric property, the following neural network’s pretrained parameters could be adapted properly to unseen datasets distribution. Third of all, train and test Hi-C interaction datasets differ but neural network can smooth such differences since interaction distributions are alike. When we generalize over distinct cell populations, the variation in the data is due to sparsity. When we generalize in terms of distinct restriction enzymes, the dissimilarity in the data is due to different Hi-C experimental biases. When we generalize in terms of distinct resolutions, dissimilarity is due to the distinct coarseness of the data. Even though all these dissimilarities exist, our neural network can still generalize which is deep learning’s important behaviour. This is also the reason why we utilize deep learning in this research.

As a future work, graph attention neural network’s number of advantages could be integrated for three-dimensional structure inference. For instance, graph attention neural network’s batching and parallelization qualities can be functional and applicable in higher resolution datasets such as resolution of less than 20 kb. In this case, since our method is generalizable, computational demands will also be lower on higher-resolution data since we pretrain the model on lower-resolution data. Moreover, a more complicated embedding alignment approach could be considered as well.

